# Green synthesized silver nanoparticles from *Moringa*: Potential for preventative treatment of SARS-CoV-2 contaminated water

**DOI:** 10.1101/2024.10.12.617998

**Authors:** Adebayo J. Bello, Omorilewa B. Ebunoluwa, Rukayat O. Ayorinde, Nneka Onyepeju, Joseph O. Shaibu, Adeniyi R. Adewole, Abeebat O. Adewole, Olusegun Adedeji, Ololade O. Akinnusi, Olajumoke B. Oladapo, Temitope S. Popoola, Oluwamodupe M. Arotiba, Joseph B. Minari, Luqman A. Adams, Joy Okpuzor, Mujeeb O. Shittu

## Abstract

Biogenic silver nanoparticles have been reported as good antimicrobial candidates. Thus, this study investigated the antiviral activity of silver nanoparticles synthesized against SARS-CoV-2. The silver nanoparticle was biosynthesized using leave extracts of *Moringa oleifera* (AgNPmo) and characterized using UV-Vis spectroscopy, Fourier transform infrared spectroscopy (FTIR), scanning electron microscopy, and X-ray diffractometry (XRD). The AgNPmo was first tested on clinical bacterial isolates, *Pseudomonas aeruginosa* (ATTC 154423) and *Staphylococcus aureus* (ATTC 209233), to ascertain its antimicrobial potential. In-vitro studies were also conducted to determine the cytotoxicity effect of AgNPmo on Vero cells. The efficacy and concentration of AgNPmo against SARS-CoV-2 were evaluated using a qPCR assay in a dose-dependent manner. The results demonstrated the successful biosynthesis and characterization of AgNPmo and its efficacy against the bacterial isolates. The AgNPmo showed low toxicity on the Vero cells. The IC_50_ from the cytotoxicity assay demonstrated the antiviral activity of the AgNPmo on the SARS-CoV-2 virus, leading to an increase in the Cycle threshold values, notably at 48 hours of incubation and at low concentrations. The results showed that the biogenic AgNPmo synthesized was cost-effective and showed both antimicrobial and antiviral potentials. These findings suggest that the nanoparticles could be a promising alternative for combating SARS-CoV-2, especially for water purification and preventing transmission.

## Introduction

Severe acute respiratory syndrome coronavirus-2 (SARS-CoV-2) is the causative agent of a contagious, life-threatening disease called Coronavirus-2019 (COVID-19) (1, 2). So far, SARS-CoV-2 has spread globally to all continents since the World Health Organization (WHO) proclaimed it a Public Health Emergency of International Concern (PHEIC) in January 2020, with more than 775 million illnesses and more than 7 million deaths confirmed as of May 2024 according to COVID-19 dashboard on World Health Organisation website. The challenges and impacts of the disease on human health, economy, and environment have led to a plethora of studies that are focused on establishing ways of curtailing the various transmission routes of the COVID-19 virus (3). One of the undermined transmission routes is by water contaminated with virus-infected human bodily excreta (4), and the virus is found to persist for seven days at 23°C (5). SARS-CoV-2 virus prevalence in wastewater has been documented to pose a risk for the transmission of COVID-19 (6, 7). Also, the virus has been found in the human gastrointestinal system (8), which can be shed via faeces and may subsequently find its way into water bodies, especially from medical wastewater, wastewater from cruise ships and aircraft that are not properly managed (9). This can lead to a developing problem, as water bodies or their components may serve as an overlooked medium for transmission of the SARS-CoV-2 virus and other viral infections. This study attempts to address this problem in a cost-effective way and could serve as a potential solution to eradicate SARS-CoV-2 in water bodies, therefore halting or slowing down the infection rate (10).

Silver nanoparticles (AgNPs) have become significantly important among other metal nanoparticles (NPs) due to their distinctive size, morphology, and environment-dependent properties, which are different from other materials’ bulk properties (11, 12). AgNPs have been used extensively as antimicrobial agents. Different studies have reported the properties and efficacy of these AgNPs against bacteria (13–15), fungi (16, 17), and viruses (18–20). The efficacy of AgNPs against microbes is due to the physicochemical properties of AgNPs, such as small particle size and high surface-area ratio, which enable their movement through cellular membranes to the target sites (18, 21, 22). Also, the easy movement of AgNPs into living cells causes accumulation, which can lead to toxic effects at very low concentrations (23). Hence, the biosynthesis of the nanoparticles provides eco-friendly and bio-compatible alternatives for microbial treatments.

This study focuses on the use of AgNPs biosynthesized from *Moringa oleifera* as a disinfectant for treating SARS-COV-2-infected water. *Moringa oleifera* Lam (drumstick tree), belonging to the family *Moringaceae*, is among the most useful medicinal trees in most of Asia and Africa. *Moringa oleifera* leaf extracts possess biocompounds with antioxidant and antibacterial activities (24). These compounds act against the growth of gram-positive and gram-negative bacteria (25). The biocompounds extracted from *Moringa oleifera* leaves also act as reducing and capping agents in the synthesis of silver nanoparticles (14), making them attractive antimicrobial agents with eco-friendly and low-cost properties.

Silver nanoparticles were successfully synthesized in this study using aqueous leaf extracts of the plant *Moringa oleifera,* and characterized using UV-Vis spectroscopy, Fourier transform infrared spectroscopy (FTIR), scanning electron microscopy (SEM), and X-ray diffractometry (XRD). The successful testing of the silver nanoparticles (AgNPs) was first carried out on clinical bacterial isolates to ascertain their antimicrobial properties. In this study, the AgNPmo showed a dose-and-time dependent antiviral activity against SARS-CoV-2, showing that the AgNPmo has the potential to be used for purification or treatment of SARS-CoV-2 contaminated water without exhibiting toxicity against living cells.

## Results and Discussion

The search for effective treatments and preventive measures against SARS-CoV-2 is still ongoing, and continuous progress in this search is pertinent to minimize the spread of the virus in water and water bodies (5, 26). Silver nanoparticles (AgNPs) have recently been studied as potential antiviral agents due to their unique properties (27). They have been shown to exhibit antiviral activities against several viruses, including influenza (28), HIV (29), and SARS-CoV-2 viruses (20).

Thus, this study aimed to assess the effectiveness of dose-dependent biogenic silver nanoparticles against SARS-CoV-2 in an aqueous environment over a period of time. The silver nanoparticles (AgNPmo) were synthesized using *Moringa oleifera* and characterized using UV-Vis spectroscopy, FT-IR, scanning electron microscopy (SEM), Energy-dispersive X-ray spectroscopy (EDX), and X-ray diffraction. The AgNPmo was first tested against clinical isolates of *Staphylococcus aureus* and *Pseudomonas aeruginosa* to ascertain its efficacy. Then, a cytotoxicity test was conducted on VeroE6 cells to determine the IC50 for the subsequent antiviral assay. The antiviral activity of AgNPmo against SARS-CoV-2 was conducted using qPCR assay in a dose and time-dependent manner.

### UV-visible analysis of the AgNPmo

The AgNPmo synthesis was first carried out using *Moringa oleifera* leaf extracts. Previous reports have shown that *M. oleifera* can serve as a reducing and capping agent in the synthesis of silver nanoparticles (30, 31), and the derived nanoparticles can act effectively against bacteria and viruses (14, 31, 32). In our study, the synthesis of AgNPmo was first ascertained visually by the color change of the reaction mixture, changing from yellowish to dark brown within 5 minutes. The UV-Vis spectroscopy revealed a 0.585 characteristic absorption peak at 420nm, revealing the formation of the AgNPmo (Fig. 1). The observed absorbance is typical of silver nanoparticles, which is within the range of 400-450 nm (33). The spectrum (in blue) showing a peak at 410 nm was taken after 12 months of the nanoparticle synthesis. This suggests that the AgNPmo retained its properties absorbance over a long period of time. The molecular interaction of the synthesized nanoparticles could be attributed to the surface plasmon resonance (SPR) due to the interaction of the free electrons found in metal-based synthesized nanoparticles with light energy (34).

**Fig 1.**
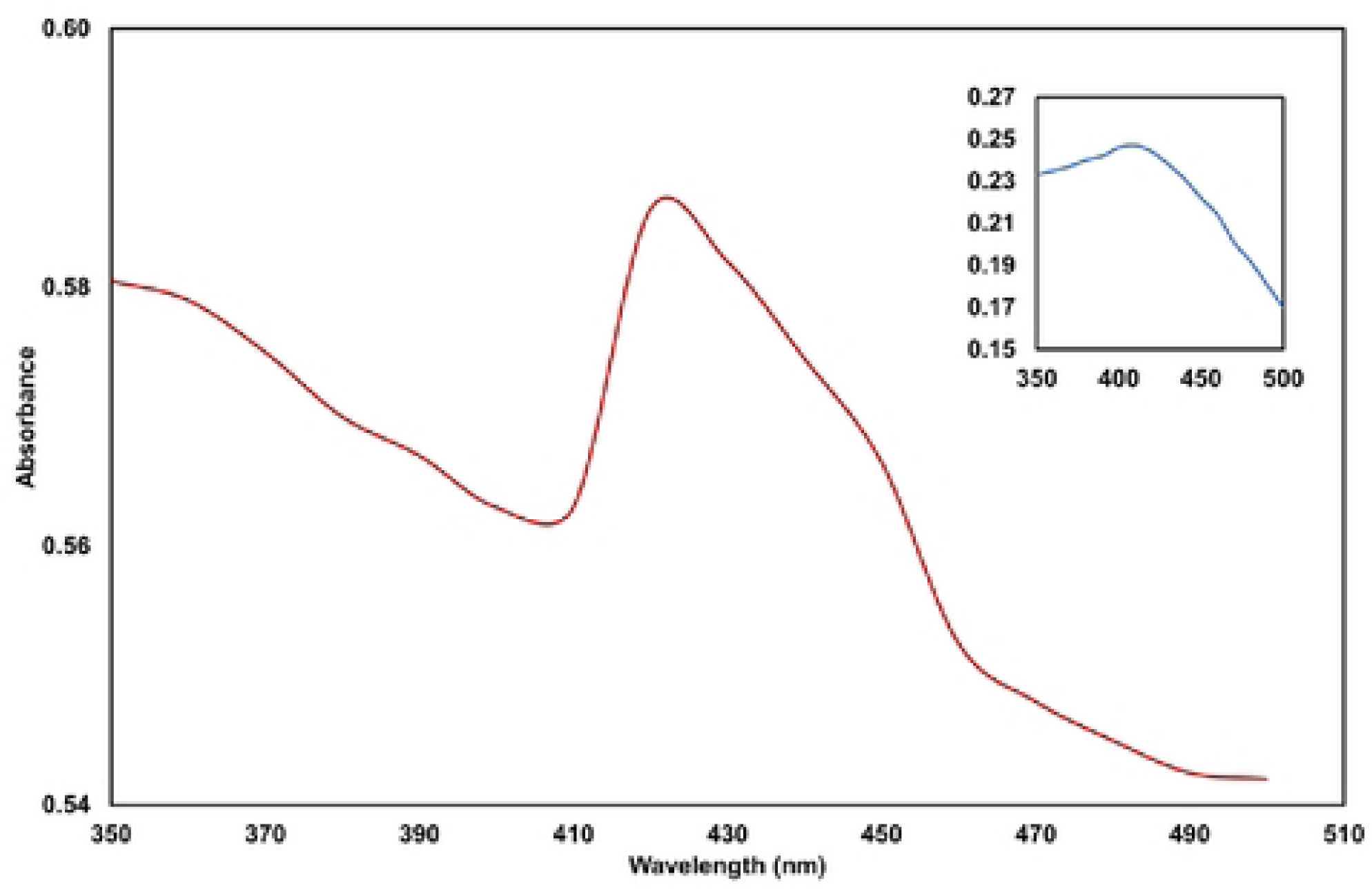
Spectrophotometry analysis of the biosynthesized silver nanoparticles Fourier transform infrared spectroscopy (FTIR) analysis of the AgNPmo.

The FTIR analysis in Fig. 2 revealed the functional groups corresponding to absorbance peaks, which are interpreted according to the spectra correlation table (35). The peaks, wavenumber (cm^-1^), and their indication are represented in Table 1. The spectra suggest that the biomolecule compounds in the *M. Oleifera* leaf extract act by reducing the Ag ions through interaction with biomolecule functional groups and thus function as capping agents in the formation of the AgNPmo within a uniform size and shape (36).

**Fig 2.**
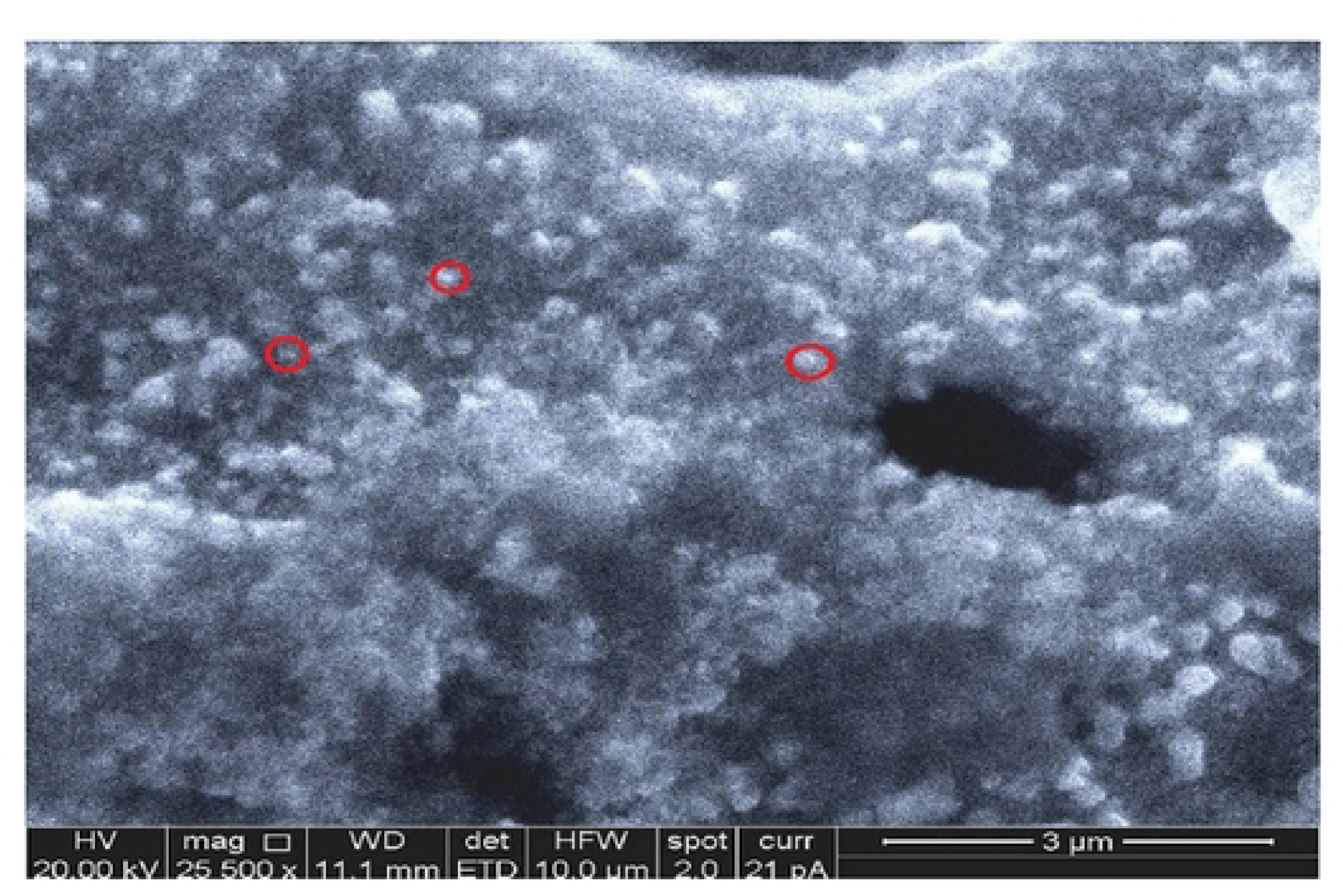
Scanning Electron Microscopy showing the particle size of the biosynthesized nanoparticles.

**Table 1.** FTIR analysis of the biosynthesized silver nanoparticles.

| Peak | Intensity | Bond/Stretching | Functional Group |
| --- | --- | --- | --- |
| 590.24 | 83.953 | C-Br stretch | Alkyl halides |
| 779.27 | 83.728 | C-Cl stretch | Amide group |
| 937.44 | 81.213 | C=H stretch | alkene |
| 1072.46 | 78.574 | C=O bend | carbonyl group |
| 1384.94 | 66.062 | C-H bend | alkane |
| 1508.38 | 81.109 | C=C bend | aromatic compound |
| 1871.01 | 89.787 | C=H stretch | aromatic compound |
| 2345.52 | 89.265 | O=C=O bend | Carbon dioxide |
| 3433.41 | 84.606 | O-H stretch | hydroxyl group |

### SEM analysis of the AgNPmo

The SEM analysis provided an inconclusive evaluation of the shape and size of the AgNPmo synthesized. Agglomeration of the nanoparticles was observed and associated with the purification procedure during synthesis. These results are similar to the findings of Moodley et al. (2018) (30). The SEM analysis identifies the morphology and particle size of the nanoparticles using the microscopy technique. In Fig 2, the AgNPmo presents an irregular surface topography with different shapes and sizes. The AgNPmo was observed at 25,000 g magnification on a 3µm scale. Most of the particles aggregated, which could result from the preparation process for SEM analysis. Some of the particles (in red circles) are less than 500nm, which may indicate that tiny nanoparticles are within the agglomerated particles. However, our SEM analysis agrees with similar reports, whereby the nanoparticles were agglomerated but retained their functionality in the antimicrobial and toxicity activities (16, 37).

### XRD analysis of the AgNPmo

The XRD analysis is a good indication of the crystalline structure and stability of nanoparticles. The AgNPmo synthesized, confirmed by the XRD pattern (Fig 3), showed diffraction spectra of 27.91°, 32.37°, and 38.30°. 44.41°, 46.20°, thus revealing the crystalline nature of the synthesized nanoparticles (38).

**Fig3.**
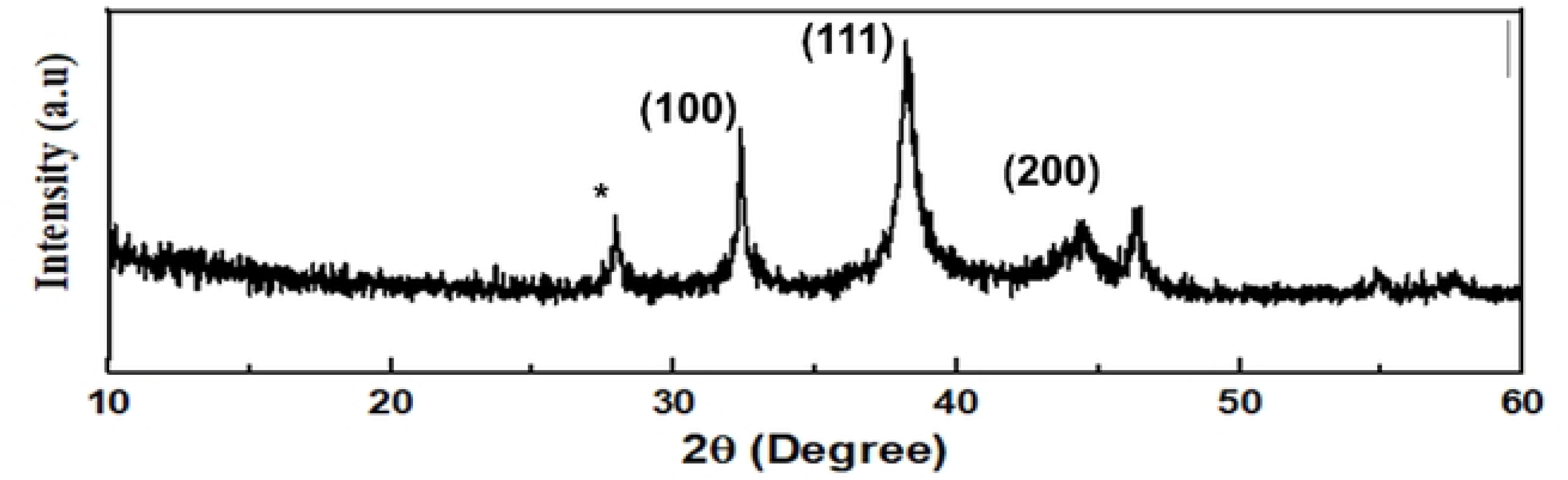
XRD analysis of the biosynthesized nanoparticles.

### Analysis of antimicrobial activity

Exposure of clinical bacterial pathogens to AgNPs has been reported to disrupt membrane permeability, prompting leakage from cells and hindering their growth and replication (39–41). First, to confirm the efficacy of the synthesized silver nanoparticles, the antibacterial activities of the AgNPmo were carried out on *Pseudomonas aeruginosa* and *Staphylococcus aureus* clinical isolates (Fig 4A and 4B). The effectiveness of AgNPs extracted from *Moringa oleifera* against the tested bacterial strains was evident in the susceptibility *of P. aeruginosa*, showing a notable sensitivity with a mean zone of inhibition (ZOI) diameter of 15.5, 11, 7.5, and 6.5 mm at AgNPmo concentrations of 100% (1.68 mg/mL), 50, 25 and 12.5% respectively (Fig 4C). Similarly, for *S. aureus,* a lesser mean ZOI diameter of 7.0, 4, 3.5, and 0 mm at similar concentrations of the AgNPmo, respectively (Fig 4D). The NC (negative control) containing DMSO without the AgNPmo showed no effect on the bacterial isolates, confirming the antimicrobial activity of the synthesized nanoparticles. The result suggests a direct correlation between the quantity of AgNPs and the expansion of the ZOI. In agreement with our study, some reports have documented the action of AgNPs against bacterial proliferation in a concentration-dependent manner (42, 43). Ultimately, AgNPs interfere with bacterial macromolecules, causing breakdown and eventual cell death.

**Fig 4.**
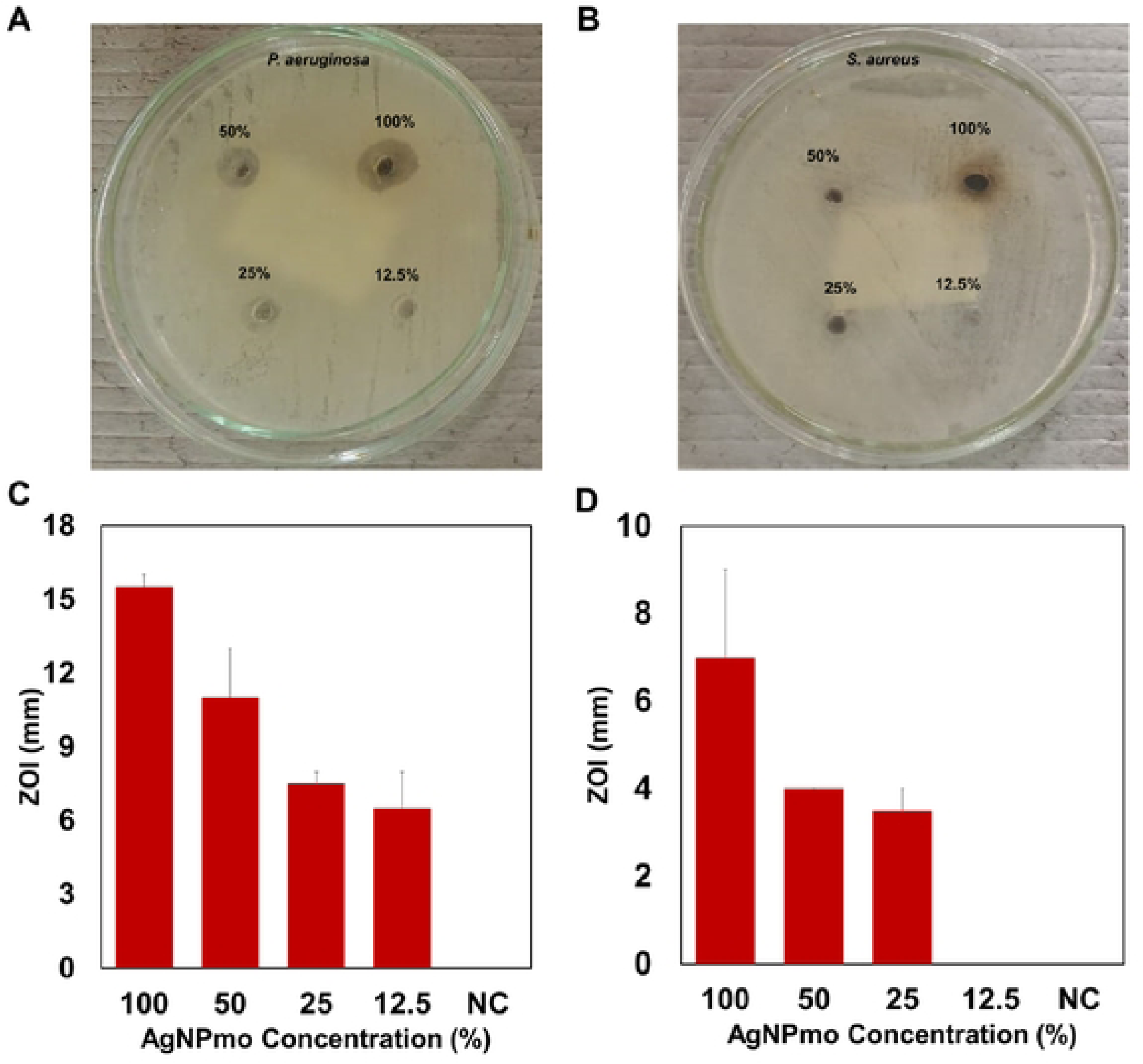
Antibacterial activities of the AgNPmo.

### Cytotoxicity assay

After confirming the efficacy of the AgNPmo by evaluating its antimicrobial activity against clinical bacterial isolates, the cytotoxicity effect on VERO cells was evaluated to determine the inhibitory concentration (IC_50_) for the subsequent antiviral study. The results revealed that the AgNPmo exhibited significant cytotoxic activities in a dose-dependent manner (Fig 5). The statistical analysis performed on data derived from the CCK-8 assay revealed IC_50_ value of 38µg/µl was determined for the 10mM stock concentration of the AgNPmo compared to the control. The viability of untreated Vero cells (control) remained unchanged at 100% cell viability. The IC_50_ value in this study is lower compared to a previous study, which reported 568g/ml IC_50_ of AgNPs biosynthesized using *Catharanthus roseus* (44). The low IC_50_ value indicates a low inhibitory effect on the viability of the Vero cells, making the AgNPmo biocompatible with human cells.

**Fig 5.**
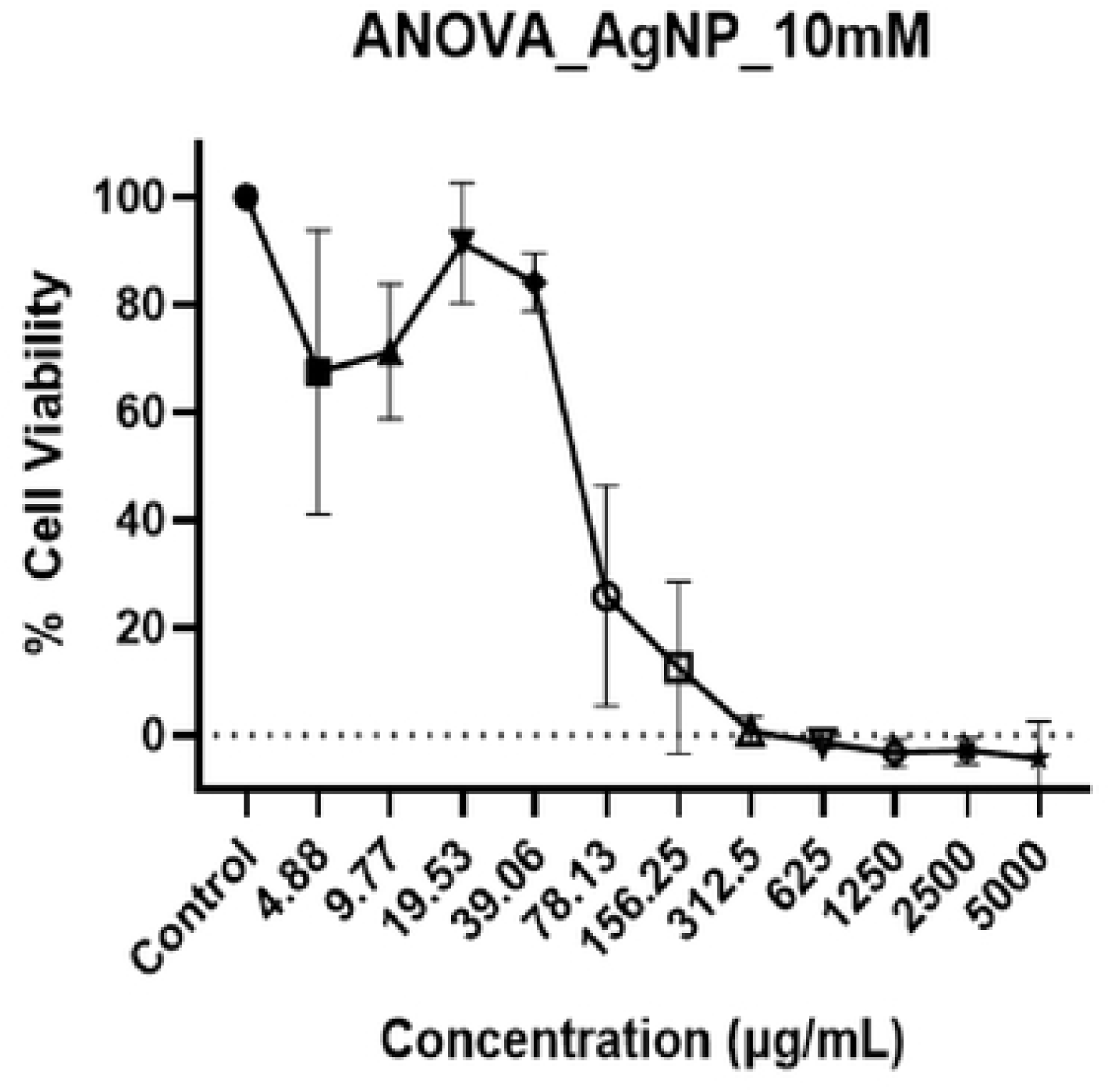
Percentage cell viability Vero cells treated with the 10mM biosynthesized silver nanoparticles.

### Quantitative RT-PCR Analysis

Quantitative PCR (qPCR) assay is a primary method of detecting the viral load of SARS-CoV-2 in various clinical specimens, including nasopharyngeal swabs stored in viral transport medium, sputum, and saliva (45). The qPCR assay targets the viral RNA, which is extracted from the clinical sample and reverse transcribed into complementary DNA (cDNA). The cDNA is then amplified using specific primers and fluorescent probes that target specific regions of the SARS-CoV-2 genome, majorly the structural proteins. The results of the qPCR for SARS-CoV-2 detection are reported as cycle threshold (Ct) values, which indicate the presence or absence of the target genes (46). The lower the Ct value, the higher the viral load in the clinical sample; a Ct value of less than 35 is considered positive for SARS-CoV-2, indicating the presence of viral RNA in the sample, while a low Ct value of less than 20 indicates a high viral load, while a high Ct value more than 30 indicates a low viral load.

This study explored the virucidal effect of biosynthesized silver nanoparticles against the SARS-CoV-2 virus at non-toxic concentrations using qPCR. To assess the dose-dependent effect of the synthesized AgNPmo on SARS-CoV-2 in a viral transport medium (VTM), the VTM was incubated with the biosynthesized AgNPmo over a period of 2 days at 24-hour intervals. The sample controls (SC) were included to confirm whether the increase in the Ct values is due to the activity of AgNPmo on the virus or environmental conditions, such as temperature, change in time of incubation, or error in the viral RNA extraction or qPCR setup.

Figs 6 and 7 represent the cycle threshold (Ct) values of the target SARS-CoV-2 genes (ORF-Lab1ab and N-genes), respectively, against the AgNPmo dilutions. Starting with the IC50 concentration from the cytotoxicity assay, the AgNPmo had a similar progressive inhibitory effect at all the concentrations for both target genes. The inhibitory effect of the AgNPmo was also time-dependent, while the Ct values increased with an increase in the hours of incubation. The Ct values for the sample controls (SC) remained steady over the incubation period. A slight upward trend was observed in the inhibitory effect of the AgNPmo with a decrease in the nanoparticle concentration for both genes (Fig 6 and 7). However, at 4.25 µg/µl concentration, there was no significant increase in the Ct values at 0 and 24 hours of incubation. The rapid increase in the inhibitory effect of AgNPmo on SARS-CoV-2 at 48 hours may be due to the long time of incubation, which could have allowed the AgNPmo to interact well with the viral particles.

**Fig 6.**
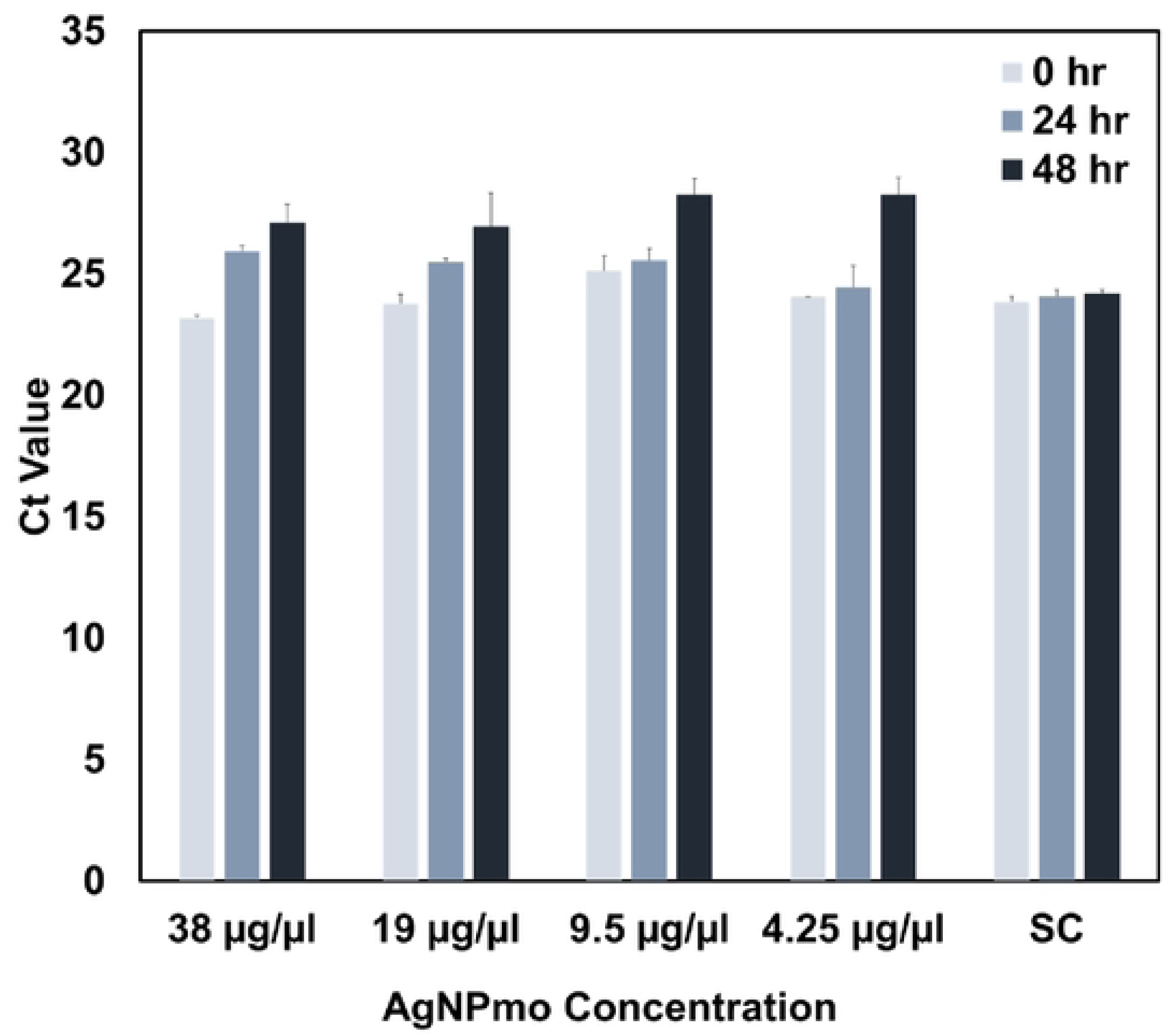
Inhibitory effect of the biosynthesized nanoparticles against SARS-CoV-2 ORF-Lab 1 gene.

**Fig 7.**
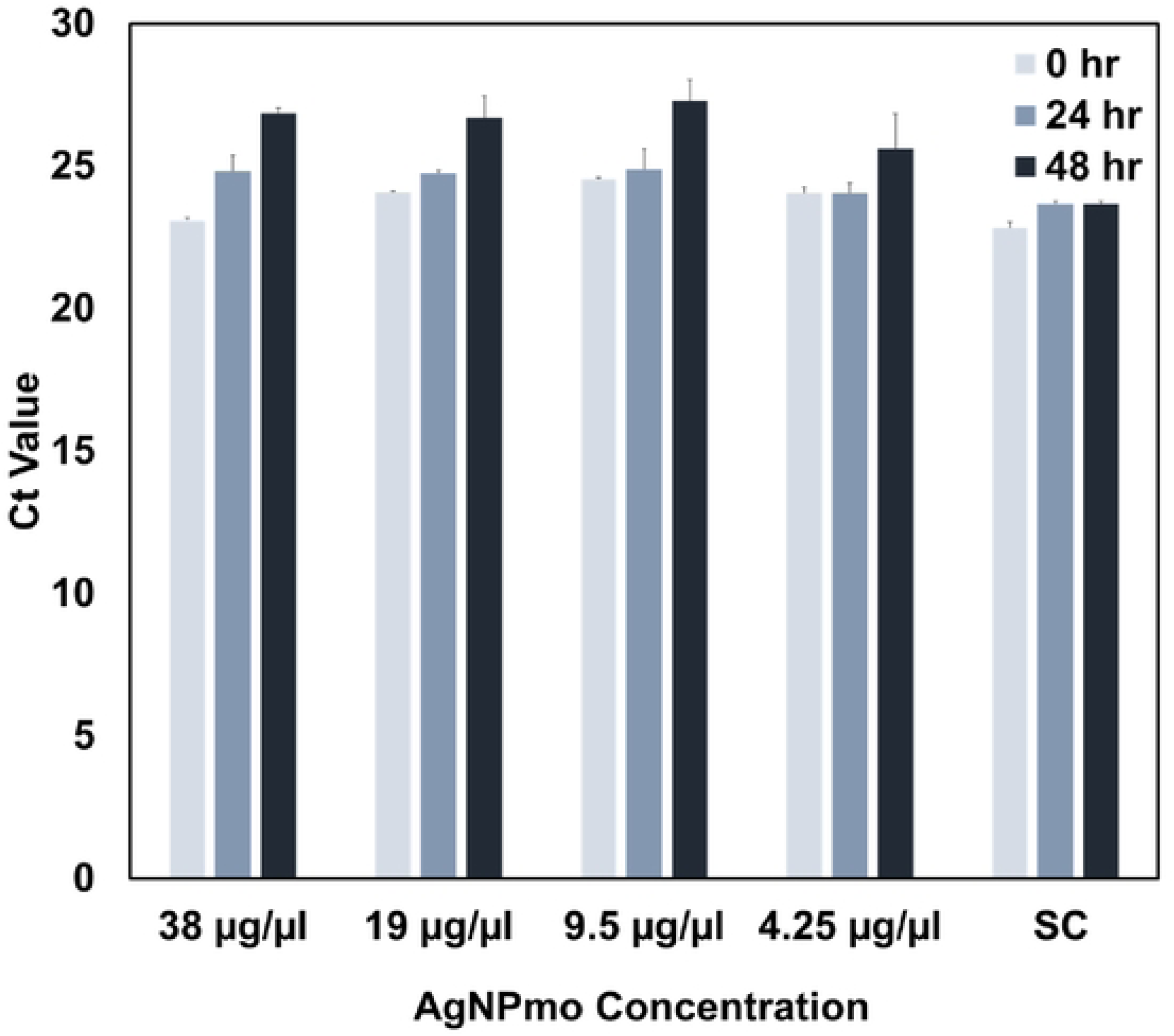
Inhibitory effect of the biosynthesized nanoparticles against SARS-CoV-2 N-gene.

The increase in Ct-values observed could be a result of the AGNPmo interacting with viral replication, thereby inhibiting the viral RNA synthesis or assembly of viral particles. The AgNPs could also have interfered with the structural proteins and inhibited their ability to bind with cell receptors or the genetic material of viruses, thereby inhibiting viral replication at the optimal concentration (47).

In summary, this study showed that the biogenic silver nanoparticles were effective in inhibiting SARS-CoV-2. The virus inside the VTM showed an increase in Ct values, indicating a reduction in its effectiveness. The results suggest that the nanoparticles are also relatively safe at controlled concentrations. The findings from this study are valuable and can be repurposed as a cost-effective and less toxic method of degrading SARS-CoV-2 particles in environmental water samples.

## Materials and methods

### Plant material and preparation of extract

Fresh *Moringa oleifera* leaves were obtained and identified at the Biological Sciences Department of the Redeemer’s University, Nigeria. The leaves were rinsed thoroughly with distilled and deionized water to remove dust particles. 20g of fresh leaf samples were submerged in 100 mL of deionized water. The mixture was heated using a microwave for 1 minute until a yellow-green coloration was observed. The crude extract was allowed to cool for 10 minutes, then filtered using Whatman® Number 1 filter paper. The filtered leaf extract was then stored in a refrigerator at 4°C for further use.

### Green synthesis of AgNPs

The silver nanoparticles were synthesized using a mixture of the aqueous leaf extract and 10 mM silver nitrate (AgNO_3_) solution prepared in deionized water. The constant microwave irradiation method was used for the nanoparticle synthesis (48). Briefly, AgNO_3_ solution and leaf extract were mixed at a ratio of 9:1, respectively. The solution was covered using a microwave-safe cling wrap and irradiated in a microwave oven operating at 400 watts for 30 seconds. The solution was allowed to stand for 5 minutes until reddish-brown coloration was observed. The overall silver nanoparticle synthesis (AgNPmo) progressed from a light yellow to a dark reddish-brown coloration.

The AgNPmo were purified by centrifugation at 1,500 rpm for 15 minutes at 4°C to avoid damaging the nanoparticles or altering their physicochemical properties. The supernatant was then discarded; the pellet was washed with deionized water, and the process was repeated. The resultant paste was dried overnight in a hot air oven at 25°C.

### Physicochemical Characterization of AgNPs

After visible coloration, the formation of the nanoparticle was first analyzed using UV-Vis spectrophotometry. The absorption spectra of the nanoparticles were analyzed within 350–500 nm wavelength using JENWAY 7305a spectrophotometer (Bibby Scientific, UK). The dried nanoparticle was diluted with distilled water, and UV-Vis spectrophotometry was repeated using nanodrop 2000 (Thermofisher, UK) after one year of synthesis. A Bruker Fourier Transform Infrared (FT-IR) spectrophotometer using the KBr pellet method was used to determine biomolecules or functional groups responsible for the reduction and capping of the prepared AgNPmo nanoparticles within the spectral range of 3500 to 500 cm^-1^.

X-ray Diffraction (XRD) analysis was performed using a D2 phaser x-ray diffractometer with a Cu Kα radiation source in the 10°-60° 2θ range, 0.02°/s scanning step size. A smooth powder surface was placed in the sample holder with the primary divergent slit confirmed as 1mm before closing the machine. The XRD machine was closed to avoid contact with the emitted radiation. The data was then transferred to the ICDD PDF-4+ software for phase analysis. Using Scanning Electron Microscopy (SEM) to estimate the dimensions of AgNPmo, the grain morphology was collected using an FEI Inspect F Scanning Electron Microscopy. Before SEM analyses, the sample was coated with carbon to make the surface conductive to enable a better signal and good image under the microscope. The SEM was performed at an accelerating voltage of 10 -15.0 kV, a spot size of 3.5 - 4.0, and a variable working distance.

### Antibacterial activity

Antibacterial activities of the AgNPs extract of *Moringa oleifera* leaves were carried out using agar well diffusion and the Minimum Inhibitory Concentration (MIC) method on *Pseudomonas aeruginosa* (ATCC 154423) and *Staphylococcus aureus* (ATCC 209233) clinical isolates. Bacteria suspension of each isolate (1.5 x 10^8^ cells/mL) was prepared and compared with 0.5 McFarland standard, which was aseptically swabbed on a Mueller Hinton Agar plate with a sterile swab stick. Diffusion wells were made using a sterile cork borer (6 mm in diameter) and placed into the agar plates. Different concentrations of the AgNPmo, 100% (1.68mg/mL), 50% (0.84mg/mL), 25% (0.42mg/mL), and 12.5% (0.21 mg/mL) prepared with 1% DMSO in two-fold serial dilution. 100 µl of each concentration was dispensed into each well labeled accordingly. The experiment was carried out in duplicate, and the plates were placed in the refrigerator for 30 minutes for the extract to diffuse into the agar. The plates were then incubated for 24h at 37 °C.

### Cytotoxicity assay

The Cell viability test of the synthesized AgNPmo was analyzed through a Cell Counting Kit-8 (CCK-8) on Vero cells. The Vero cells (2 × 105) were seeded into each well of a 96-well plate and cultured for 24 hours before any treatment. The AgNPmo treatment groups were administered serial dilutions of the stock solution from 5mg/mL to 0 mg/mL.10 µl CCK-8 reagent was added, and after 3 hours of incubation under humidified conditions at 37 °C, the color development of the samples was measured at a test wavelength of 450 nm and a reference wavelength of 630 nm using an 800 TS microplate reader (Biotek Instruments, USA). The experiment was done in duplicate.

The relative cell viability was calculated as follows:

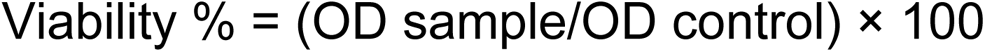

The half maximal inhibitory concentration (IC50) was calculated using the statistics software Graph Prism version 9.

### Quantitative RT-PCR Assay for antiviral activity

The COVID-19-positive samples preserved inside the Viral Transport Medium (VTM) were used for the experiment. The samples used were obtained from the Nigeria Institute of Medical Research (NIMR), and the ethical approval (IRB/23/029) was obtained from the ethical committee of the institute reviewer board. AgNPmo stock concentration was diluted serially with sterile VTM to make 38, 19, 9.5, and 4.75 µg/µl based on the IC_50_ report from the cytotoxicity assay. 100 µl of each dilution was mixed with 100 µl SARS-CoV-2 contaminated VTM and incubated for two days (0, 24, and 48 hours) at 25°C with period shaking. Viral RNA was extracted using the Qiagen RNA nucleic acid extraction kit (Qiagen, Hilden, Germany) at 0-, 24-, and 48-hour intervals. 60µl of the viral RNA was eluted after dry spinning the column for 2 minutes.

The quantitative RT-PCR was performed using the SCODA SARS-CoV-2 Fast PCR assay protocol according to Shaibu et al. (2023) (49). The final reaction mixture of 25µl contained 7µl of the SCODA reagent A, 13µl of the SCODA reagent B, and 5µl of each sample. PCR amplification was achieved using a QuantStudio™ 5 Real-Time PCR System (ThermoFisher Scientific, UK) at optimized PCR conditions of 52°C for 10 minutes, 95°C for 10 seconds, 95°C for 5 seconds (40 cycles) and 56°C for 30 seconds (40 cycles). The reaction was performed in duplicate. Three controls were set up: First, the sample control (SC) contained the reaction mixture without AgNPmo. The negative control (NC) contained the reaction mixture and RNAse-free water without the sample. The third control contained the qPCR kit positive control (PC), replacing the sample. Both the NC and PC are used to control the quality of the PCR process.

### Statistical analysis

Arithmetic means and standard error (SE) were calculated for the antimicrobial and antiviral experiment using the Microsoft Excel 2016 version.

